# Benchmark of lncRNA Quantification for RNA-Seq of Cancer Samples

**DOI:** 10.1101/241869

**Authors:** Hong Zheng, Kevin Brennan, Mikel Hernaez, Olivier Gevaert

## Abstract

Long non-coding RNAs (lncRNAs) emerge as important regulators of various biological processes. Many lncRNAs with tumor-suppressor or oncogenic functions in cancer have been discovered. While many studies have exploited public resources such as RNA-Seq data in The Cancer Genome Atlas (TCGA) to study lncRNAs in cancer, it is crucial to choose the optimal method for accurate expression quantification of lncRNAs. In this benchmarking study, we compared the performance of pseudoalignment methods Kallisto and Salmon, and alignment-based methods HTSeq, featureCounts, and RSEM, in lncRNA quantification, by applying them to a simulated RNA-Seq dataset and a pan-cancer RNA-Seq dataset from TCGA. We observed that full transcriptome annotation, including both protein coding and noncoding RNAs, greatly improves the specificity of lncRNA expression quantification. Pseudoalignment-based methods detect more lncRNAs than alignment-based methods and correlate highly with simulated ground truth. On the contrary, alignment-based methods tend to underestimate lncRNA expression or even fail to capture lncRNA signal in the ground truth. These underestimated genes include cancer-relevant lncRNAs such as *TERC* and *ZEB2-AS1*. Overall, 10–16% of lncRNAs can be detected in the samples, with antisense and lincRNAs the two most abundant categories. A higher proportion of antisense RNAs are detected than lincRNAs. Moreover, among the expressed lncRNAs, more antisense RNAs are discordant from ground truth than lincRNAs when measured by alignment-based methods, indicating that antisense RNAs are more susceptible to mis-quantification. In addition, the lncRNAs with fewer transcripts, less than three exons, and lower sequence uniqueness tend to be more discordant. In summary, pseudoalignment methods Kallisto or Salmon in combination with the full transcriptome annotation is our recommended strategy for RNA-Seq analysis for lncRNAs.

**AUTHOR SUMMARY:** Long non-coding RNAs (lncRNAs) emerge as important regulators of various biological processes. Our benchmarking work on both simulated RNA-Seq dataset and pan-cancer dataset provides timely and useful recommendations for wide research community who are studying lncRNAs, especially for those who are exploring public resources such as TCGA RNA-Seq data. We demonstrate that using full transcriptome annotation in RNA-Seq analysis is strongly recommended as it greatly improves the specificity of lncRNA quantification. What’s more, pseudoalignment methods Kallisto and Salmon outperform alignment-based methods in lncRNA quantification. It is worth noting that the default workflow for TCGA RNA-Seq data stored in Genomic Data Commons (GDC) data portal uses HTSeq, an alignment-based method. Thus, reanalyzing the data might be considered when checking gene expression in TCGA datasets. In summary, pseudoalignment methods Kallisto or Salmon in combination with full transcriptome annotation is our recommended strategy for RNA-Seq analysis for lncRNAs.

## INTRODUCTION

Long non-coding RNAs (lncRNAs) are a diverse class of RNA molecules that are more than 200 nucleotides in length and do not encode proteins [1]. While functional classification is lacking for most lncRNAs, based on their genomic proximity to protein-coding genes and the direction of transcription, lncRNA are often classified into antisense, intronic, bidirectional, intergenic, or overlapping RNAs [1]. GENCODE, the database that provides the largest set of standardized lncRNA annotations, defines over 15,000 human lncRNA genes in the current release (release 27, https://www.gencodegenes.org). Compared with protein-coding genes, lncRNAs are shorter, lower-expressed, less evolutionarily conserved, and expressed in a more tissue-specific manner [2]. lncRNAs have recently emerged as an essential class of regulatory elements for many biological processes including imprinting, cell differentiation, and development [3]. They are often disrupted in human diseases including cancer [4]. They may interact with DNA, RNA, and proteins, and exert regulatory roles through a variety of mechanisms. Based on their molecular functions, lncRNA may act as i) signals, which are indicators of transcriptional activity; ii) decoys, which bind to and titrate away protein targets such as transcription factors; iii) guides, which direct regulatory complexes or transcription factors to specific targets and regulate gene expression in *cis* or *trans*, and iv) scaffolds, which serve as central platforms where relevant molecular components in cells are assembled [5].

lncRNAs have been shown to be important in the pathogenesis of human diseases, especially in cancer, and many cancer-relevant lncRNAs have been identified [6, 7]. For example, Hox transcript antisense RNA (*HOTAIR*), one of the most well-characterized lncRNAs, promotes breast cancer metastasis through recruitment of Polycomb chromatin remodeling complex to silence the *HOXD* gene cluster [8]. *HOTAIR* is overexpressed in breast, liver, pancreatic, lung, and pancreatic cancers [9]. *CDKN2B-AS1*, an antisense lncRNA encoded by the *CDKN2B* locus, epigenetically silences nearby tumor suppresser genes and promotes oncogenesis. [10]. Telomerase RNA component (*TERC*), the critical RNA component of telomerase polymerase, serves as a template for the enzyme telomerase reverse transcriptase (TERT) to elongate telomeres. Variants and copy-number changes at the TERC locus have been associated with cancer risk and progression [6]. The lincRNA *MEG3* inhibits cell proliferation by downregulating MDM2 and promoting p53 accumulation [11].

Discovery of oncogenic and tumor suppressor lncRNAs has led to an increased interest in investigation of lncRNAs as potential cancer drug targets and biomarkers. Hence, it is critical to accurately determine lncRNA expression in cancer research. RNA sequencing (RNA-Seq) has been widely used for massive-parallel gene expression quantification. Many lncRNAs are transcribed by RNA polymerase II and are often 5'-capped, spliced and polyadenylated [12]. In a study of an earlier GENCODE version, it was estimated that 39% of lncRNAs contains poly(A) motifs, compared to 51% for protein-coding transcripts [2]. Thus, although RNA-Seq was originally designed to inspect protein-coding genes expression, a significant proportion of lncRNAs can also be examined within RNA-Seq data derived from samples that have been prepared even using poly (A) enrichment method. Multiple tools for processing RNA-Seq data have been developed in recent years. While some studies have benchmarked RNA-Seq analysis workflows [13, 14], their focus has been primarily on protein-coding genes. Moreover, there is no accepted gold standard pipeline yet which method performs best to quantify expression of lncRNAs. There have been many studies that explore lncRNA expression profile in cancer using publically available RNA-Seq datasets such as those generated by The Cancer Genome Atlas (TCGA), which provide a rich source of lncRNA expression data in large cancer patient populations. Among those studies, the analysis of the lncRNA expression profile of breast cancer samples in TCGA revealed different subtypes of breast cancer and subtype-specific over-expression of HOTAIR [15]. Also, the analysis of 13 cancer types in TCGA revealed highly cancer site-specific lncRNA expression and dysregulation [16]. As the interest in studying lncRNAs in cancer grows, it is necessary to determine which algorithms perform best in lncRNA expression quantification, as it is of uttermost importance to understand the differences and limitations of each of them. and to follow the best practice of RNA-Seq analysis. Because of the lower expression and different properties of lncRNAs to protein coding genes, we hypothesized that the processing and analysis RNA-Seq data for lncRNA expression may be subjected to different technical biases and challenges, and that special considerations may be necessary to optimize the pipeline specifically for lncRNAs.

To investigate the performance of different methods on the quantification of lncRNAs and find out what types of lncRNAs can be discovered by the mainstream RNA-Seq methods, we applied five popular methods, Kallisto [17], Salmon [18], RSEM [19], HTSeq [20], and featureCounts [21] on both simulated and real datasets. Kallisto and Salmon are so-called pseudoalignment methods, as they do not align reads to the reference genome; instead, they assign reads to a set of compatible transcripts. The alignment-free feature makes pseudoalignment methods much faster than alignment-based methods, since the latter requires alignment of the sequencing reads to the genome or transcriptome. While pseudoalignment methods are widely accepted to improve the efficiency of protein-coding genes expression analysis, their performance for lncRNAs detection has not been evaluated. In this analysis, we used STAR [22] for alignment of the reads to both the genome and the transcriptome, before applying the alignment-based methods RSEM, HTSeq, and featureCounts.

## RESULTS

### Full transcriptome annotation improves the specificity of RNA quantification

We used RSEM [19] to simulate RNA sequencing reads based on 63 randomly selected TCGA samples from 21 cancer types. To mimic noise in real data, 5% noise was introduced to the simulated samples. To evaluate the effect that different transcriptome annotations has on the quantification of gene expression, we built three transcriptome annotation sets, 1) full annotation with all 58,037 genes in GENCODE release 25 (**Supplementary Table 1**), 2) 19,950 protein-coding genes 3) 14,203 lncRNAs. Gene quantification was performed using each of the tools and with each of three annotation sets as the transcriptome reference. Figure 1 shows that using only a subset of annotation over-estimates gene expression, especially for lncRNAs genes. When using full annotation, the percentage of expressed lncRNAs detected by Kallisto and Salmon is about 10%, which is very close to the ground truth, whilst when using only the lncRNAs for transcriptome annotation, the percentage of expressed lncRNAs rises to 50%, indicating that many reads have been falsely assigned to lncRNA genes. The influence of incomplete transcriptome annotation is less drastic for protein-coding genes, but there is still a slight increase of the percentage of expressed genes when using only protein-coding reference, compared to full annotation (Figure 1B). Thus, full annotation improves the specificity of RNA quantification; therefore, it was used in the following analysis.

**Figure 1.**
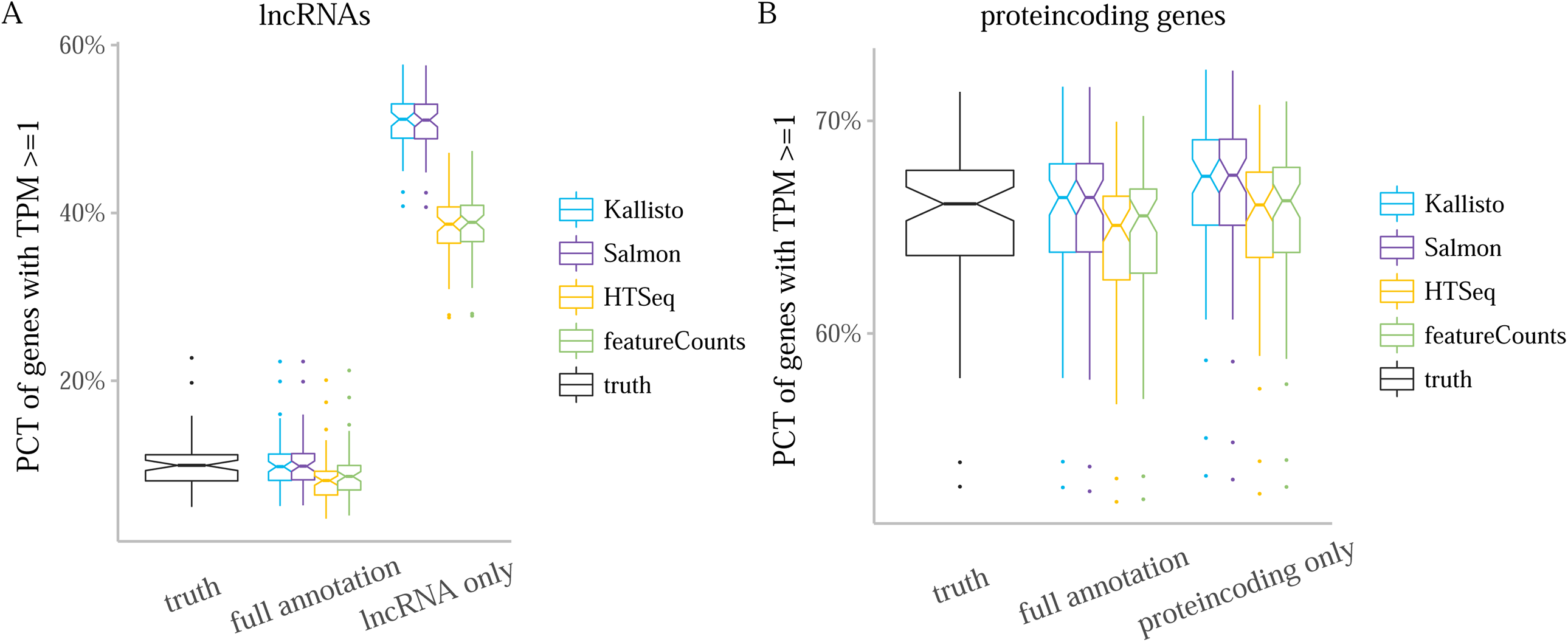
Effect of full annotation and lncRNA (protein-coding) only annotation. Boxplot of percentages of (**A**) lncRNAs and (**B**) protein-coding genes with TPM>=1 detected with each tool, using full annotation or lncRNA/protein-coding annotation. The ground truth expression used for simulation was also plotted for comparison. Each point in the boxplot represents one sample. PCT, percentage.

### Pseudoalignment methods outperform alignment-based methods for lncRNA expression quantification

Pseudoalignment methods detect expression of more genes than alignment-based methods across the simulated datasets (Figure 2A). The ‘ground truth’ percentages of annotated genes that are expressed in the simulated dataset (average across all samples) is 10.2 ± 3.1% (mean ± sd), 65.5 ± 3.6%, and 27.2 ± 2.0%, for lncRNA, protein-coding, and all genes, respectively. The percentage of expressed lncRNAs detected by Kallisto and Salmon is 10% on average, while HTSeq and featureCounts detect 8-9% of lncRNAs on average (**Supplementary Table 2**).

**Figure 2.**
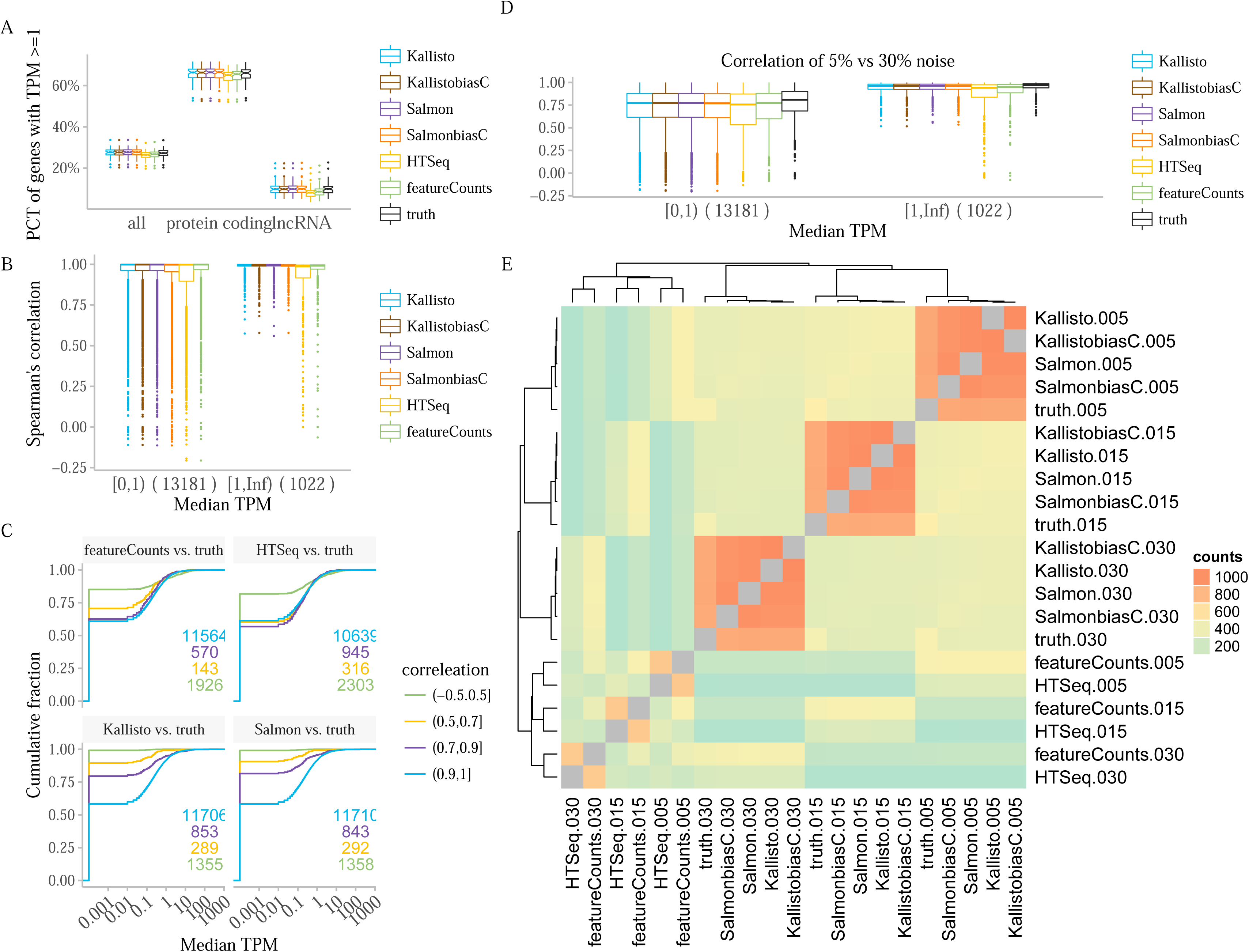
Pseudoalignment methods outperform on simulated datasets. **A)** Box plot of percentage of expressed genes detected with each tool. Each point in the boxplot represents one sample. **B)** Spearman correlation of lncRNA TPM values of simulated samples reported by each tool compared with ground truth. Each point in the boxplot represents one lncRNA gene, categorized by two groups according to median TPM values. Spearman correlation was calculated across 63 samples for each gene. **C)** Cumulative fraction of TPM values from ground truth in different tiers of correlation (comparing each method and ground truth). The labelled number indicates how many genes belong to a certain correlation tier. **D)** The Spearman correlation of TPM between 5% and 30% noise levels. Each point in the boxplot represent one lncRNA. Spearman correlation was calculated across 63 samples. **E)** Similarity matrix of combinations of tools and noise levels. Each grid in the matrix is the number of times that two entries were clustered in the same group, which is counted from hierarchical clustering of 1,022 expressed lncRNAs. The group number for cutting the hierarchical clustering dendrogram was set as four. In both single gene and method level clustering, Euclidean distance and average linkage were used. PCT, percentage; biasC, bias correction option of the method was used.

Gene expression estimates from pseudoalignment methods are more concordant (i.e. correlated better, see Methods for the definitions) with ground truth than alignment-based methods (Figure 2B). Besides, genes with higher expression levels tend to be more concordant with the ground truth. Specifically, there are 1,022 expressed lncRNAs with median TPM above one across all samples in the ground truth. Only two of these genes are discordant (see Methods for the definitions) for pseudoalignment methods, while for HTSeq and featureCounts there are 187 and 143 discordant genes, respectively (**Supplementary Figure 1A**). For Kallisto and Salmon, almost all lncRNAs whose correlation with ground truth is below 0.5 are lowly-expressed and more genes in higher correlation tiers are expressed. On the other hand, for HTSeq and featureCounts, gene expression level is not clearly associated with the correlation with ground truth (Figure 2C). To investigate the effect of different noise levels, another two sets of ground truths were generated using the same TCGA samples, with 15% and 30% noise introduced into the reads. Since simulated data is based on the TPM values from real samples, read counts for higher noise levels are lower. However, the effect of different noise levels on the percentage of expressed genes measured by TPM values is subtle (**Supplementary Figure 2A and B**). Due to random fluctuations in the simulation process, the TPM values of the ground truths at different noise levels vary slightly, but are highly correlated for genes with higher expression (median Spearman’s correlation 0.97 for expressed genes). However, HTSeq and featureCounts show a higher degree of variability between different noise levels, suggesting that they are more sensitive to noise. (Figure 2D).

For each of the 1,022 expressed lncRNA genes, hierarchical clustering was performed across all combinations of noise levels and tools to evaluate the similarity of each method’s measurement to ground truth (Figure 2E). Kallisto and Salmon are often clustered together with the ground truth under different noise levels. In addition, although featureCounts and HTSeq form another cluster, these two alignment-based methods are less similar to each other than the two pseudoalignment methods are to each other. Finally, the two genes that are discordant in all methods compared with ground truth are shown in **Supplementary Figure 1B**. Alignment-based methods fail to detect the two genes in the samples, regardless of noise levels, while pseudoalignment methods tend to over-estimate gene expression in some samples.

### Pseudoalignment methods estimate lncRNA expression more concordantly on real pan-cancer data

A total of 210 TCGA samples from 21 cancer types were examined with Kallisto, Salmon, HTSeq, featureCounts, and RSEM. Similar to simulated datasets, pseudoalignment methods detect a higher ratio of lncRNAs compared with alignment-based methods (Figure 3A). The mean percentage of annotated lncRNAs detected as expressed by Kallisto and Salmon is 16% However, only 13% of lncRNAs were detected with featureCounts and HTSeq, while RSEM only detects 10% of lncRNAs. Therefore, alignment-based methods detect fewer expressed lncRNAs in real data, which, based on our simulated data analysis, likely represents an underestimation. Next, in terms of correlation between methods, pseudoalignment methods behave much more concordantly than alignment-based methods (Figure 3B). Among the 1,346 expressed lncRNAs, only one gene is poorly correlated between Kallisto and Salmon (Spearman correlation less than 0.7). However, there are 85 to 234 expressed lncRNAs that are poorly correlated between pseudoalignment and alignment-based methods, and even between the alignment-based methods themselves. Similarly, cumulative TPM in different correlation tiers showed that genes with higher correlation between methods tend to be expressed at a higher percentage. The majority of lncRNAs are highly-correlated between Kallisto and Salmon (**Supplementary Figure 3**). For each of the 1,346 expressed lncRNAs, hierarchical clustering was performed across the tools (Figure 3C). Kallisto and Salmon are always grouped together (1,335/1346 times), and so are HTSeq and featureCounts (1,097/1,346 times).

**Figure 3.**
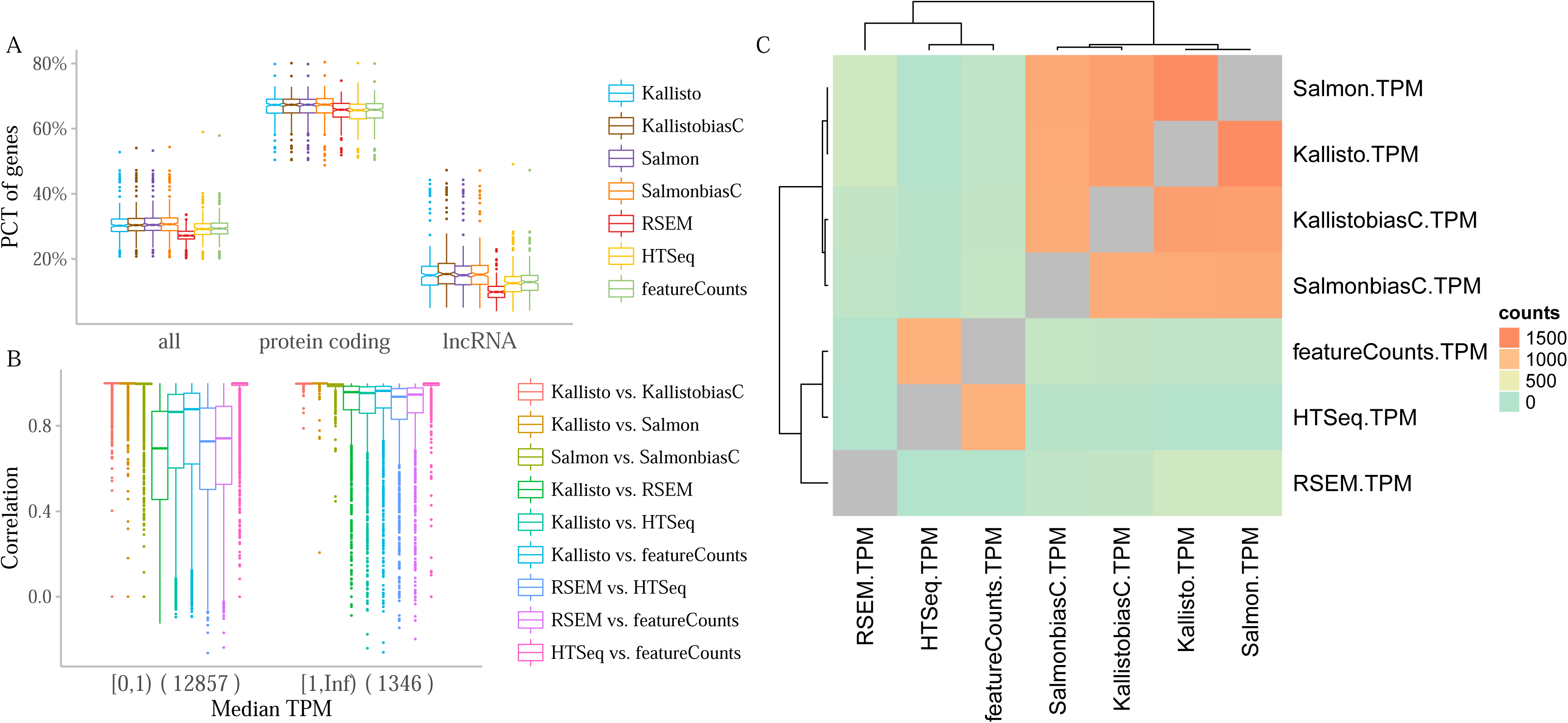
Pseudoalignment methods perform more consistently on pan-cancer datasets. **A**) Box plot of percentage of expressed genes detected with each tool. Each point in the boxplot represents one sample. **B)** Spearman correlation of TPM values of lncRNAs, comparing two methods. Each point in the boxplot represents one lncRNA, categorized by two groups according to median TPM values. **C)** Similarity matrix of different methods. Each grid in the matrix is the number of times that two methods were clustered in the same group, which is counted from clustering of 1,346 expressed lncRNAs. The group number for cutting the hierarchical clustering dendrogram were set as three. In both single gene and method level clustering, Euclidean distance and average linkage were used. PCT, percentage; biasC, bias correction option of the method was used.

### Properties of expressed and discordant lncRNAs

Out of the 14,203 annotated lncRNAs in the GENCODE v25, 53.1% are lincRNAs and 38.9% are antisense. However, in the analyzed pan-cancer dataset, out of the 1,346 expressed lncRNAs, 36.5% are lincRNAs and 53.9% are antisense (Figure 4A), indicating that a greater proportion of antisense lncRNAs are detected than lincRNAs (chi-squared test p-value < 2.2e-16). The other 9.7% lncRNAs are sense intronic, sense overlapping, and 3 prime overlapping lncRNAs. Over three quarters of lncRNAs have only one transcript, and only 6.6% of them are expressed, while over 18% of lncRNAs with two transcripts or more are expressed. Hence, lncRNAs with more transcripts are expressed at a higher percentage (Figure 4B). In addition, 16% of lncRNAs with only one exon are expressed, which is 1.7-2.2 times higher than lncRNAs with more exons (Figure 4C).

**Figure 4.**
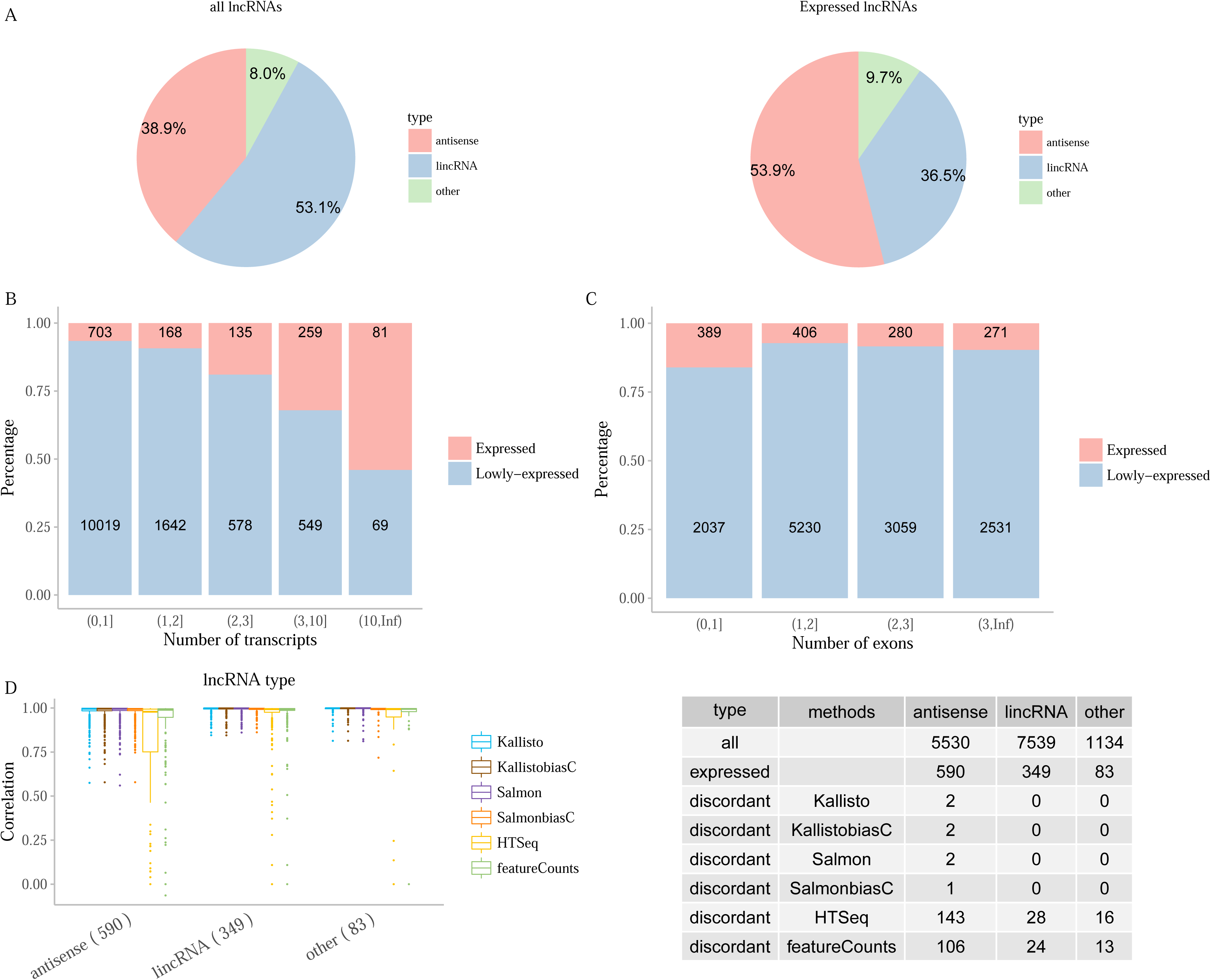
Features of expressed and discordant lncRNAs. **A**) The categories of 14,203 lncRNAs in GENCODE release 25 and 1,346 expressed lncRNAs in pan-cancer dataset. **B**) The number of transcripts and **C**) the number of exons of lncRNAs. Expressed, lncRNAs with median TPM above one across pan-cancer dataset. Lowly-expressed, lncRNAs with median TPM below one across pan-cancer dataset. **D)** lncRNA type of 1,022 expressed lncRNAs in simulated dataset. Spearman correlation was calculated comparing each method and ground truth. Numbers in brackets in x-axis labels are the number of genes in certain category. Tables on the right show the number of discordant lncRNAs for each method.

There are 1,022 expressed lncRNAs in the simulated datasets. Antisense RNAs are the major source of discordant lncRNAs. Around 70% of the discordant lncRNAs for alignment-based methods, including the two discordant lncRNAs for all methods, are antisense. About 18-24% of expressed antisense lncRNAs are discordant, while only 7-8% of expressed lincRNA are discordant, indicating that antisense lncRNAs are more susceptible to mis-quantification from alignment-based methods (chi-squared test p-value < 1e-04) (Figure 4D). The lncRNAs with fewer transcripts tend to be more discordant. Specifically, when measured by alignment-based methods, over 20% of expressed lncRNAs with only one transcript are discordant, which is more than two times higher compared to those with two transcripts or more (**Supplementary Figure 4A**). In terms of the number of exons, lncRNAs with two exons have the highest ratio of discordance (**Supplementary Figure 4B**). The fraction of unique kmers (31mers) per lncRNA, i.e., the degree of ‘uniqueness’, has a greater effect on the performance of alignment-based methods. For lncRNAs with low uniqueness (less than 20% unique sequences,) the mean correlation for HTSeq and featureCounts with ground truth is below 0.3, and the correlation increases as the uniqueness of the lncRNAs increases. Kallisto and Salmon perform well even for lncRNAs with very low uniqueness. However, uniqueness only explains part of the discordance, as there are still considerable number of discordant genes even with high uniqueness (**Supplementary Figure 4C**).

### Examples of concordant and discordant lncRNAs

Several lncRNAs that were previously linked to cancer are shown in Figure 5. Some of them are expressed only in a subset of samples, such as *TERC* and *HOTAIR*. HTSeq and featureCounts often underestimate RNA expression, as expected. For *HOTAIR, NKILA*, and *CCAT1* on the right panel, all the methods perform equally well and cluster with corresponding ground truth. However, lncRNAs on the left panel are not profiled accurately. HTSeq and featureCounts often form distinct cluster away from ground truth and the rest of the methods. Moreover, in two examples all the ground truth in different noise levels cluster together, indicating that even pseudoalignment methods do not always profile lncRNAs accurately. More discordant lncRNAs examples are shown in **Supplementary Figure 5** [16]. HTSeq and featureCounts again fail to capture or underestimate these lncRNAs. For some lncRNAs, such as *ENSG00000176868.2*, none of the methods captures the signal as in the ground truth in some samples.

**Figure 5.**
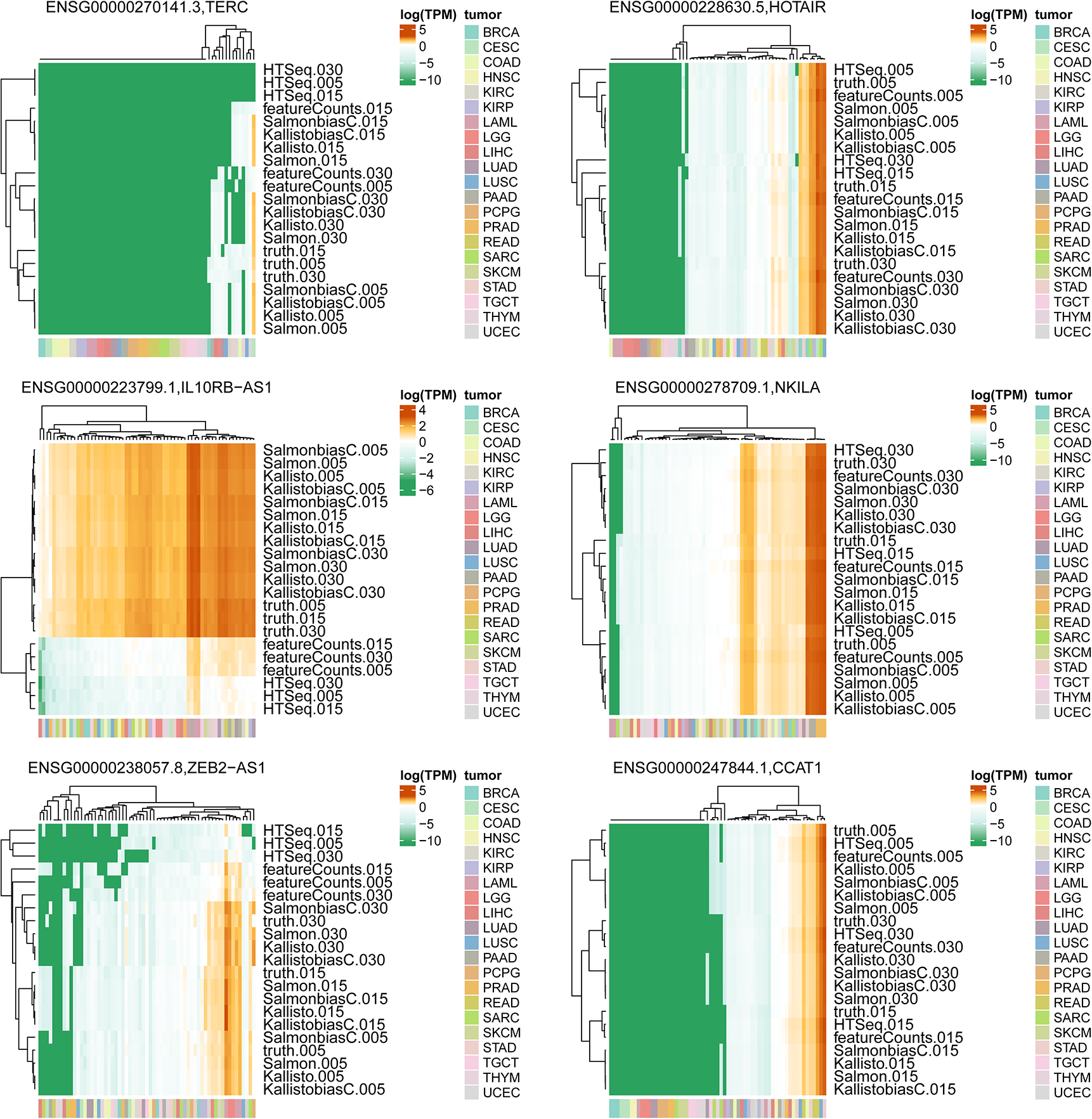
Examples of lncRNAs in cancer. The lncRNAs that were previously reported to play a role in cancer are shown in the simulated dataset. Heatmap shows the TPM (log transformed) of each method under different noise levels, across 63 samples. Euclidean distance and average linkage were used. 005, 5% noise level; 015, 15% noise level; 030, 30% noise level; biasC, bias correction option of the method was used.

Next, we investigated cancer-site specific lncRNAs. The number of cancer-site specific lncRNAs varies among different cancer types (**Supplementary Figure 6**). A total of 1,381 lncRNAs were found to be specifically expressed in the acute myeloid leukemia (LAML) samples analyzed in this study, which is much higher than other cancer sites. Lower grade glioma (LGG) has 121 cancer-site specific lncRNAs. The other cancer types, pheochromocytoma and paraganglioma (PCPG), testicular germ cell tumor (TGCT), prostate adenocarcinoma (PRAD), and liver hepatocellular carcinoma (LIHC), have 20-41 cancer-site specific lncRNAs.

## DISCUSSION

Although most publically available RNA-Seq datasets such as TCGA mainly capture protein-coding genes and they may not be optimal for lncRNAs profiling, a fair amount (over 10% on average) of lncRNAs can still be detected (TPM above one) in the samples in this study. The RNA of TCGA samples were captured by poly(A) enrichment methods, thus, ideally the lncRNAs with poly(A) tails can be characterized from the data, if they are expressed. These public resources provide rich opportunities for studying the expression and function of lncRNAs in cancer in a cost-effective way. It is thus critical to choose the right method for accurate expression quantification of lncRNAs.

In this benchmarking work, the major expressed lncRNA categories detected are antisense and lincRNAs. However, other less abundant categories, sense intronic, sense overlapping, and 3 prime overlapping ncRNAs, can also be detected. Among the expressed lncRNAs, over half of them are antisense RNAs, although they only account for 36% of all lncRNAs in the GENCODE annotation. Antisense lncRNAs were once considered as transcriptional noise, but later were shown to influence gene expression from transcription and translation to RNA degradation [23]. The abundantly expressed antisense RNAs in cancer samples are worth examining more closely in future studies. However, it is worth noting that antisense RNAs contribute to the majority of discordant lncRNAs measured by alignment-based methods, which needs extra attention if the antisense RNAs of interest are quantified using these methods in future research.

The two pseudoalignment methods, Kallisto and Salmon, perform highly concordantly in both the simulated dataset and the pan-cancer dataset, as previously reported [24, 25]. Compared to alignment-based method, they detect more lncRNAs in each sample, which is similar to the level of ground truth in the simulated datasets. They are also highly consistent with the ground truth at gene-level comparisons, with only two highly-expressed lncRNAs that are discordant between the measured TPM and the ground truth. Kallisto is faster than Salmon, and utilizes slightly less memory (**Supplementary Figure 7**), but they are comparable in terms of accuracy. The bias correction option of these two tools takes slightly longer time, but we didn’t observe noticeable difference in terms of concordance with ground truth whether using the bias correction option or not. On the contrary, the alignment-based method HTSeq and featureCounts, have a much higher number of discordant genes. Kallisto and Salmon always cluster together with ground truth, while HTSeq and featureCounts always form another cluster. It is worth noting that HTSeq is the default workflow for TCGA data stored in GDC data portal. Since it is not the optimal quantification method, re-analyzing the data might be considered when checking gene expression in TCGA datasets.

The discordant genes for HTSeq and featureCounts are mainly i) antisense RNAs, ii) lncRNAs with only one transcript, and iii) lncRNAs with less than three exons. HTSeq and featureCounts tend to underestimate the abundance of genes, including some of the well-known lncRNAs in cancer, such as *TERC* and *ZEB2-AS1* (Figure 5). Furthermore, out of the 1,022 genes with TPM above one in the simulated dataset, about 119 lncRNAs do not have any reads assigned to them by HTSeq and featureCounts. A subset of the underestimated genes can be explained by the low sequence uniqueness, such that the tools discard many of the ambiguously mapped reads by default.

Since the simulated data is generated using RSEM, we cannot fairly compare RSEM in the simulated data experiments. However, in the real pan-cancer dataset, RSEM underperforms when comparing to pseudoalignment methods. It detects the lowest percentage of expressed lncRNAs and there are over 150 expressed lncRNAs that are discordant between RSEM and other methods. It also does not cluster with HTSeq and featureCounts, but forms a distinctive cluster in single gene level clustering. Thus, the pseudoalignment methods Kallisto and Salmon are recommended over the three alignment-based methods investigated.

Although for most of the analysis Kallisto and Salmon perform better than alignment-based methods and cluster with ground truth, on certain occasions pseudoalignment methods are also suboptimal, as in the case of some cancer-relevant lncRNAs, including *TERC, IL10RB-AS1, ZEB2-AS1* (Figure 5), and examples in **Supplementary Figure 5**. In recent studies, *IL10RB-AS1* gene is shown to be differentially expressed in diffuse large B cell lymphoma [26]. Upregulation of *ZEB2-AS1* is suggested to promotes tumor metastasis and predicts poor prognosis in hepatocellular carcinoma [27]. Under such occasions, although pseudoalignment methods do not accurately quantify lncRNA expression in all the samples, they still perform better than alignment-based methods.

Next, lncRNAs are expressed in a cell type-, tissue-, developmental stage- or disease state-specific manner [12], which partly explains that not many lncRNAs have median TPM above one across the cancer types. In the samples examined in this study, cancer-site specificity is also observed. There are 1,670 cancer-site specific lncRNAs in total, with the majority associated with LAML. The drastic difference on the numbers of cancer-site specific lncRNAs between LAML and all other types is likely due to the difference between lymphatic cells vs. solid tissues. The cancer-site/tissue specificity of expressed lncRNAs implies the importance of choosing the right method to capture the subtle signal.

Lastly, the comparison in this analysis was performed at the gene-level. If the goal is to examine transcript-level expression, HTSeq and featureCounts are not suitable for the purpose. They are developed explicitly for gene-level read counts. When they count reads mapped to transcripts rather than genes, reads mapped to exons shared by several transcripts will then be considered ambiguous and discarded by default. Therefore, for transcript-level analysis, Kallisto, Salmon, or RSEM should be used.

In summary, considering the consistency with ground truth, flexibility with both genes and transcripts analysis, and the running time (**Supplementary Table 3**), pseudoalignment methods Kallisto and Salmon in combination with the full transcriptome annotation including protein coding genes, lncRNAs, and others, is the recommended strategy for RNA-Seq analysis for lncRNAs.

## MATERIAL AND METHODS

### Definitions

We would like to clarify or define relevant terms. i) Genes and transcripts: a transcript, sometimes also referred to as an isoform, is composed by exons. A gene is a collection of transcripts. Transcripts of the same gene often share overlapping exons. In this analysis comparison was performed at the gene-level. ii) Expressed genes: when describing a single sample, “expressed genes” refer to genes with TPM (transcript per million) value equal or above one in that particular sample. When describing multiple samples, “expressed genes” refer to the genes with median TPM value equal or above one across the cohort of samples. iii) Concordant genes: these are defined as genes whose Spearman correlation of TPM values with ground truth is more than 0.7, when compared across a cohort of samples. iv) Discordant genes: these are defined as genes whose Spearman correlation of TPM values with ground truth is less than 0.7, when compared across a cohort of samples.

### Datasets

210 samples from 21 cancer types in TCGA were randomly selected and downloaded from Genomic Data Commons (GDC) Data Portal. These RNA-Seq samples were prepared with poly(A) selection method and non-strand-specific protocol. Simulation of RNA-Seq reads under different noise levels (0.05, 0.15, 0.3) was performed with the RSEM command *rsem-simulate-reads*, using the model information and quantification results of 63 TCGA samples from 21 cancer types. Reads QC were performed with Trim_galore [28], with the setting “-q 15 --stringency 3 --length 15 -q 15 --paired”.

### Reference genome and transcriptome

Human transcriptome version GENCODE release 25 (GTF file and transcriptome fasta file) were downloaded from GENCODE FTP site. The GENCODE release 25 collects 58,037 genes and 198,093 transcripts, among them there are 19,950 protein-coding genes (**Supplementary Table 1**). If defining lncRNA as non-coding, 3-prime overlapping ncRNA, antisense, bidirectional promoter lncRNA, long intergenic noncoding RNAs (lincRNA), macro lncRNA, sense intronic, and sense overlapping, there are 14,203 lncRNAs. Compared to protein-coding genes, lncRNAs genes are shorter, have fewer transcripts and fewer exons.

The primary assembly of human genome GRCh38 was also downloaded from the same site. The index for Kallisto and Salmon was built using the transcriptome fasta file, while the index for RSEM and STAR was built using transcriptome GTF file and GRCh38 genome sequences.

Gene features (number of transcripts, number of exons, transcript length) were generated from GTF file using in-house script. Unique kmers of genes were generated using script from Computational Genomics Analysis and Training (CGAT).

### Pipelines

Four pipelines were applied to process the datasets. All the reads were unstranded RNA-Seq data, and the relevant parameters were set for each tool.

Kallisto, version 0.43.0, quant mode. Default parameters were applied. The ‘--bias’ option was also added when correcting for sequence bias. Stand-specific option: default.

Salmon, version 0.8.2, quant mode. Default parameters were applied. The ‘-seqBias –gcBias’ option was also added when correcting for sequence and GC content bias. Stand-specific option: default or ‘-l IU’. Kallisto and Salmon measure the expression level of each transcript by default. To get gene-level expression results, the package tximport [29] was used.

RSEM, version 1.3.0. Reads were first mapped to human transcriptome with STAR (version 2.5.3a, two-pass mode). The command ‘*rsem-calculate-expression’* of RSEM was then used for transcript and gene level quantification. Stand-specific option: ‘--forward-prob 0.5’.

HTSeq, version 0.7.2. Reads were first mapped to human genome with STAR (version 2.5.3a, two-pass mode). Fragments were then counted for each gene according to GTF file. Stand-specific option: ‘-s no’.

featureCounts, version 1.5.2. Reads were first mapped to human genome with STAR (version 2.5.3a). Fragments were then counted for each gene according to GTF file. Stand-specific option: ‘-s 0’. HTSeq and featureCounts output read count for each gene. TPM values were generated from read counts with in-house scripts.

To compare the CPU time consumed per sample for each tool, 21 samples were chosen at random and processed with each tool in a single CPU thread.

### Clustering and heatmaps

Hierarchical clustering was performed for the sample-method matrix. Euclidean distance and average linkage were used for both the columns and rows. The clustering dendrogram was cut into 3-4 groups and the number of times that every two tools are clustered together were counted and used to construct a similarity matrix.

### Cancer-site specific lncRNAs

Cancer-site specific lncRNAs in this study are defined as i) median TPM above one in samples from one cancer site and below one in other cancer samples; ii) maximum log TPM in other cancer samples less than one; iii) p values from Wilcoxon signed-rank test between the specific cancer and other cancer samples less than 0.0001. Only primary tumor samples and leukemia samples are included in the analysis.

## AVAILABILITY

All the scripts and dockerized tools were put in GitHub (zhengh42/RNASeq_pipeline and zhengh42/Dockerfiles).

## ACKNOWLEDGEMENT

This work was supported by National Institute of Dental & Craniofacial Research (NIDCR) (U01 DE025188), the National Institute of Biomedical Imaging and Bioengineering (R01 EB020527), and the National Cancer Institute (U01 CA217851), all of the National Institutes of Health. The content is solely the responsibility of the authors and does not necessarily represent the official views of the National Institutes of Health. The funders did not play a role in the design and execution of this study.

## SUPPLEMENTARY DATA

**Supplementary Figure 1 Discordant genes between truth and each method**

**A)** Venn diagram of expressed discordant genes. Genes are defined as discordant if Spearman’s correlation of TPM with ground truth is less than 0.7. Spearman correlation was calculated using 63 samples with 5% noise levels. **B)** Heatmap shows the TPM (log transformed) of each method under different noise levels, across 63 samples. The two genes that are discordant in all methods are shown.

**Supplementary Figure 2 Percentage of expressed genes across all noise levels**

**A)** The percentages of genes with read count above 10 detected with each tool was shown. The higher the noise level, the lower the percentage of genes with read count above 10. **B)** The percentages of genes with TPM above one detected with each tool remain unchanged in different noise levels. Each point in the boxplot represent one sample in the simulated dataset.

**Supplementary Figure 3 Cumulative fraction of TPM values in different tiers of correlation in pan-cancer dataset**

The labelled number indicates how many genes belong to certain correlation tier. Median TPM was calculated based on TPM values from the two methods in comparison. Spearman correlation was calculated across 210 samples.

**Supplementary Figure 4 Features of expressed lncRNAs in simulated dataset.**

A) Number of transcripts, B) Number of exons, and C) unique kmers of 1,022 expressed lncRNAs in simulated dataset. Spearman correlation was calculated comparing each method and ground truth. Numbers in brackets in x-axis labels are the number of genes in certain category. Tables on the right show the number of discordant lncRNAs for each method.

**Supplementary Figure 5 Examples of discordant lncRNAs in cancer.**

The lncRNAs that were previously reported to have copy number changes in cancer are shown in the simulated dataset. Heatmap shows the TPM (log transformed) of each method under different noise levels, across 63 samples. Euclidean distance and average linkage were used. 005, 5% noise level; 015, 15% noise level; 030, 30% noise level; biasC, bias correction option of the method was used.

**Supplementary Figure 6 Cancer-site specific lncRNAs lncRNAs in pan-cancer dataset**

**A)** Number and **B**) Heatmap of cancer-site specific lncRNAs. Cancer-site specific lncRNAs are defined as those with median TPM above one in the specific cancer, median TPM below one in other cancer samples, maximum log TPM below one in other cancer samples, and p values from Wilcoxon signed-rank test between groups less than 0.0001. TPM values in this plot were reported by Kallisto. COAD and READ are considered from the same cancer site. Only 184 primary tumor samples and leukemia samples are included in the analysis.

**Supplementary Figure 7 CPU time for each tool**

CPU time was calculated with 21 samples. Single thread was used. For STAR, RSEM, HTSeq, and featureCounts, default parameters were applied. For Kallisto and Salmon, bootstrap (100 times) or bias correction options were also added. bias, bootstrap (+) and bias correction (+); nobias, bootstrap (+) and bias correction (−); noboot_bias, bootstrap (−) and bias correction (+); noboot_nobias, bootstrap (−) and bias correction (−).

**Supplementary Table 1 Genes and transcripts in GENCODE release 25**

**Supplementary Table 2 Wilcoxon signed-rank test of percentages of expressed genes detected by each tool, compared with truth data**

**Supplementary Table 3 Summary of the characteristic and performance of tools.**

The tools suitable for profiling the expression of genes, transcripts, and exons were marked. ^1^The tools output transcript-level quantification by default. Gene-level expression is aggregated from transcript-level result using the tximport package. ^2^The time of STAR processing was added.

